# Cryo-EM structural studies of the agonist complexed human TRPV4 ion-channel reveals novel structural rearrangements resulting in an open-conformation

**DOI:** 10.1101/2020.10.13.334797

**Authors:** Mathieu Botte, Alexander K. C. Ulrich, Ricardo Adaixo, David Gnutt, Andreas Brockmann, Denis Bucher, Mohamed Chami, Nicolas Bocquet, Ulrich Ebbinghaus-Kintscher, Vera Puetter, Andreas Becker, Ursula Egner, Henning Stahlberg, Michael Hennig, Simon J. Holton

**Author notes:** A.U.: Proteros biostructures GmbH, Martinsried, Germany, V.P., U.E., A.B. and S.J.H.: NUVISAN Innovation Campus Berlin GmbH, Germany.

## Abstract

The human transient receptor potential vanilloid 4 (hTRPV4) ion channel plays a critical role in a variety of biological processes. Whilst the activation of hTRPV4 gating properties has been reported for a broad spectrum of stimuli, including synthetic 4α-phorbols, the molecular basis of the activation is poorly understood. Here we report the novel cryo-EM structure of the hTRPV4 determined in the presence of the archetypical phorbol acid agonist, 4α-PDD. Complementary mutagenesis experiments support the EM-identified binding site as well as allowing rationalization of disruptive mutants located outside of the 4α-PDD binding site. This work represents the first structural information of hTRPV4 in a ligand-induced open conformation. Together, our data reveal the underlying molecular mechanisms resulting in the opening of the central pore and ion-channel activation and provide a structural template for designing inhibitors targeting the open-state conformation of hTRPV4.

## INTRODUCTION

The transient receptor potential (TRP) ion channel superfamily is involved in a broad range of physiological processes and their dysregulation results in various diseases [1–3]. TRP channels, formed from homo- or hetero-tetramers, contain a central pore that functions as a cation channel. Mammalian TRP family proteins have been classified into six subfamilies based on sequence similarity [2]. The vanilloid TRP subfamily (TRPV1-6) has been further classified into two subgroups; the thermosensitive, Ca^2+^ non-selective TRPV1-4 channels (known as thermo-TRPs) and TRPV5-6, which are both insensitive to temperature and Ca^2+^ selective [4–8].

TRPV4, first described as an osmotically activated channel [9, 10], is a thermo TRP ion channel that has been shown to play a prominent role in a multitude of biological processes and dysregulation of its activity has been associated with several human diseases [11–13]. Various studies have identified potential for therapeutic intervention in a range of pathologies including pain, gastrointestinal, neurodegenerative disorders, cancer and lung diseases, including most recently COVID-19 [13–17]. This extensive range of biological roles reflects the broad diversity of TRPV4 modulators that include temperature, endogenous ligands or lipids and synthetic agonists and antagonists [17–20].

In response to these diverse signals, TRP channels can adopt either a closed, non-conducting or an open, ion-conducting state. In addition to these two extreme states - open or closed - the channels are thought to undergo frequent transitions to additional, intermediate states, for example inactive, partially and transiently closed conformations [21]. Structures of TRPV family members in different functional states have provided insights into the structural elements and conformational changes involved in gating mechanisms. Much of this understanding was initially derived from TRPV1 structures in open, closed and a partially-activated state [22–24]. Subsequently determined structures of other TRPV family members have identified gating mechanisms that are broadly conserved across the TRPV family as well as revealing specific mechanisms utilized by single or sub-family members. Key structural elements involved in the gating mechanism include the pore helix between the helices S5 and S6, the helix S6 itself, the S4-S5 linker and the amphipathic TRP helix. The *Xenopus tropicalis* TRPV4 (xTRPV4) structure revealed selected structural elements adopt unique conformations not previously observed in other TRPV channels [25]. For example, the S1-S4 bundle and S5-S6 pore domains are much closer than has been observed in other TRP channels and the outer pore is unusually wide and only accommodates a single ion -binding site. This conformation may represent an inactive nonconductive state that is structurally different to the resting-closed state observed in other TRP closed conformation structures [26]. Based on these features, it has been postulated that TRPV4 may display different gating behavior compared to other TRPV channels [25, 26].

Despite the wealth of TRPV channel structural information generated in recent years, only a very limited number of open-conformation ligand-complexed TRPV structures have been reported. Consequently, the molecular mechanisms through which many of the different stimuli influence the TRPV gating mechanism are not fully understood. Complex structures in the presence of either endogenous lipids or exogenous ligands have identified two hot-spots, or binding sites, within the transmembrane domain (TMD) region, through which channel gating may be modulated. The first of these sites – the vanilloid binding site – is located between the S3 helix, the S4-S5 linker and the S6 helix of the adjacent subunit [22–24]. Binding of an endogenous lipid to this site in the mammalian TRPV1, TRPV2 and TRPV3 proteins promotes and stabilizes the closed conformation of the ion-channel [27]. Lipid displacement, for example through direct competition with the RTX or capsaicin agonists, leads to a switch to the open TRPV1 ion-channel conformation. The second site – the voltage-sensing like domain (VSLD) binding site - is located at the interface between the cytoplasmic side of the S1-S4 helices and the membrane-facing side of the TRP helix. As with the vanilloid binding site, lipids or effector molecules binding at the VSLD site have been observed to regulate the ion-channel conformation and activity [28]. Intriguingly, whilst TRPV3 adapts an active open conformation upon binding of 2-APB at this site, the TRPV6 channel adopts an inactive closed-conformation upon 2-APB binding [29–31]. These contrasting effects upon binding of the same molecule to analogous binding sites in different proteins highlights the versatility and sensitivity of ion channel responses to external stimuli.

The association of hTRPV4 with various diseases has motivated research towards the identification of activity modulators. Reported agonists include 4α-phorbol 12,13-didecanoate (4α-PDD), a synthetic phorbol ester tool compound (EC_50_ 200 nM) [11, 32] and GSK1016790A (EC_50_ 2 nM) [33]. Whilst site-directed mutagenesis experiments have identified several residues that disrupt the 4α-PDD mediated activation of hTRPV4 [32, 34, 35], the molecular mode-of-action has remained elusive.

In this contribution we report the first high-resolution structural data for the hTRPV4 ion channel. In the presence of the weak hTRPV4 agonist 4α-PDD, the structure reveals for the first time the open-state conformation. Through comparison with models of the closed state human protein, we have gained insights into the mechanism of 4α-PDD mediated hTRPV4 ion-channel activation.

## RESULTS

### Determination of 4α-PDD bound hTRPV4 structure

To study the molecular basis of 4α-PDD mediated activation of the hTRPV4, we expressed and purified a truncated form of the protein which included all conserved TRPV4 sequence segments including the TMD, TRP, linker and ARD domains. The truncated construct, encompassing residues 148-787, resulted in significantly improved insect-cell recombinant protein expression levels compared to the full-length construct (871 residues).

We next tested the pharmacological response of the truncated and full-length hTRPV4 constructs upon GSK1016790A stimulation in both transfected oocytes and insect cell lines. Both full-length and truncated channels could be functionally reconstituted in transfected oocytes and generated currents of up to 30μA in response to GSK1016790A stimulation (Fig. 1). Similarly, GSK1016790A induced channel activation was observed for both constructs following transient expression in insect cells (Extended Data Fig. 1). Whilst the truncated hTRPV4 construct retained its ion permeability (after stimulation), it was expressed less efficiently and displayed an approximate 10-fold increase in the EC_50_ for the channel activation by GSK1016790A compared to the full-length channel (Extended Data Fig. 1). Similar effects have been reported for other similarly truncated thermostable TRP channels [25, 36, 37]. Taken together, these observations support the use of the truncated hTRPV4 construct to characterize molecular activation mechanisms of the full-length native hTRPV4 channel.

**Fig. 1.**
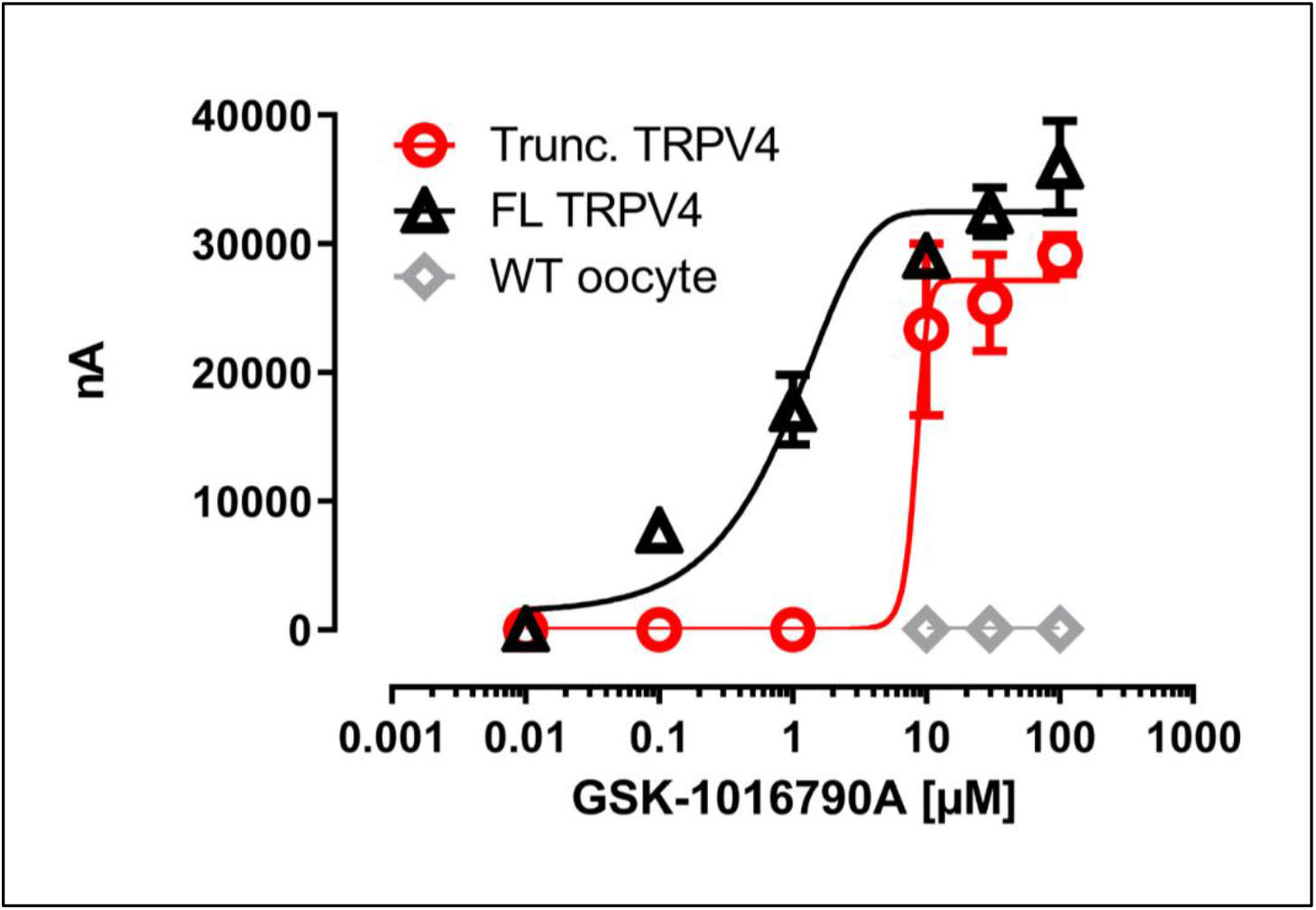
Electrophysiological characterization of full-length and truncated hTRPV4 channels. GSK1016790A dose-response curves for oocytes expressing either full-length (black) or truncated (red) cryo-EM hTRPV4 channel construct. The two-electrode voltage-clamp recordings were performed at a holding potential of −60mV. Half maximal effective concentration (EC_50_) for full-length (EC_50_ 0.83μM) and truncated hTRPV4 (EC_50_ 8.43μM) channels were calculated from logarithmic fitting of the data. Each data point represents the average of 3-12 independent measurements.

**Extended Data Fig. 1.**
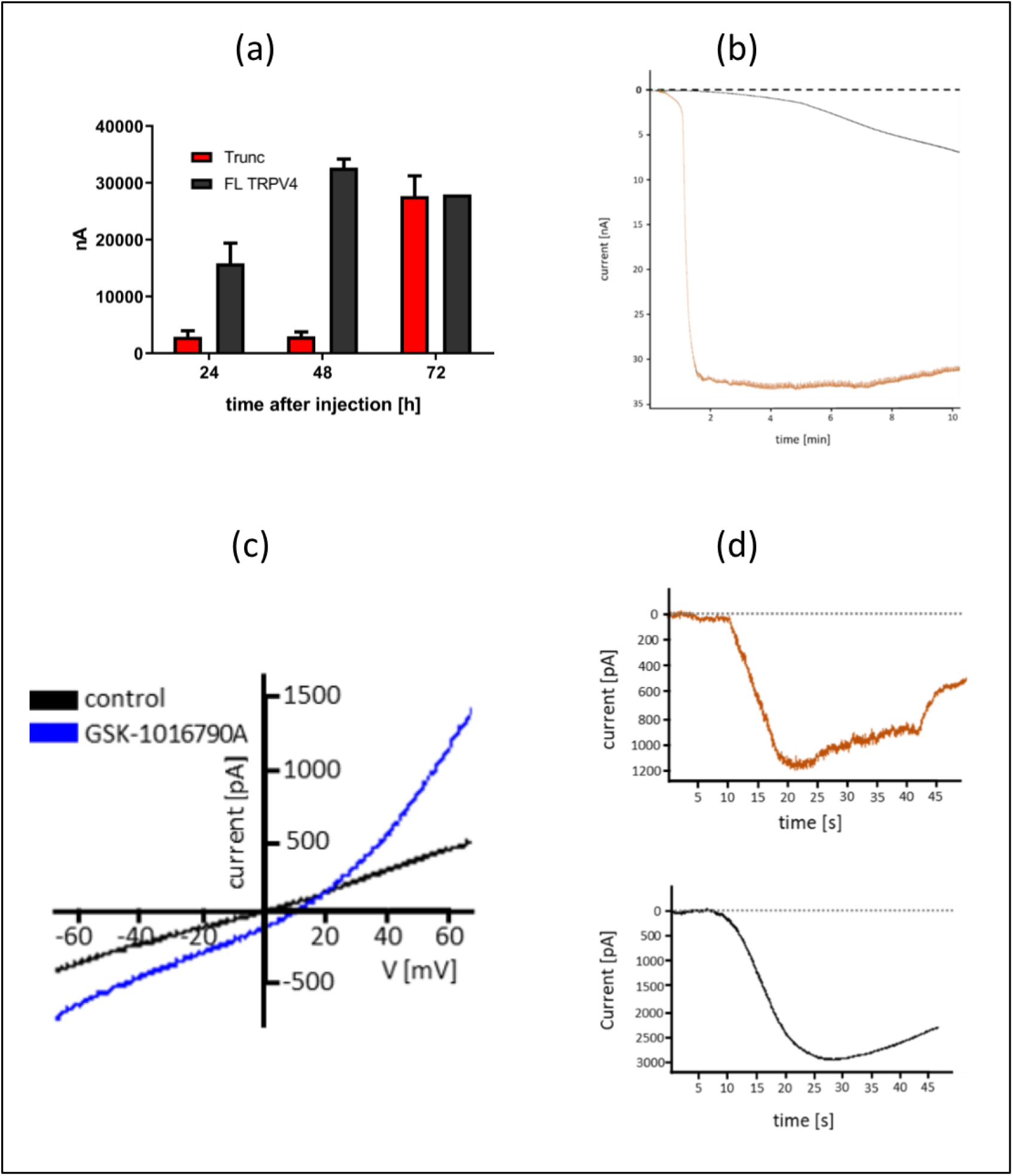
Full-length and truncated electrophysiological hTRPV4 characterization. (**a**) Expression levels of hTRPV4 in oocytes. Two electrode voltage-clamp (TEVC) measurements showing the time-dependent maximal current induced by high concentrations of GSK1016790A after RNA injection for full-length TRPV4 (black) and truncated TRPV4 (gray) constructs. In these measurements the cells were clamped at a holding potential of −60 mV. The full-length TRPV4 channel reached its strongest signal after 48h and was stable for at least 72h. Comparable currents were observed for the truncated construct after 72h. Based on these observations dose-response curves shown in **Fig. 1** were determined 48h and 72h after RNA injection for full-length and truncated TRPV4, respectively. (**b**) Representative oocyte two-electrode voltage clamp (TEVC) responses for hTRPV4 variants stimulated with GSK1016790A and 4α-PDD. Representative traces showing the time-dependent current responses for oocytes expressing either truncated TRPV4 treated with 30μM GSK1016790A (blue) or full-length TRPV4 treated with 30μM 4α-PDD (gray). In the measurements the cells were clamped at a holding potential of −60 mV. The truncated hTRPV4 channel exhibited a fast response to GSK1016790A that reached a steady-state current with the first 60s. In contrast, the full-length hTRPV4 construct exhibited a slower response to 4α-PDD. In all 9 trials a continuously slow increase in the current was observed throughout the course of the experiment (9 minutes). The truncated TRPV4 construct did not respond at the concentrations of 4α-PDD tested. Based on the reduced GSK1016790A efficacy in activating the truncated channel (**Fig. 1**) it is likely that the concentrations of 4α-PDD required to activate the truncated channel are not technically achievable in this assay system. (**c**) Characteristic I/V curves obtained by voltage ramps in presence and absence of GSK1016790A. The traces showing the voltage current relationship of High-Five cells expressing the full-length hTRPV4 treated with 0.1 μM GSK1016790A (blue) or untreated (black). The voltage ramps started from holding potential −70 mV to +70 mV within 70 ms. GSK1016790A exposure started 3 s prior to the measurement. (**d**) Representative whole cell voltage clamp responses of TRPV4 variants to GSK1016790A. Representative whole cell voltage clamp responses of high five insect cells transiently expressing either full-length hTRPV4 channel (lower trace) or truncated hTRPV4 channel (upper trace). The full-length channel responded to both 0.3μm (shown) and 1μM (not shown) GSK1016790A with currents of 2065 ± 220 nA (n=8 of 11 tested cells). A significant truncated hTRPV4 response was also observed upon stimulation with 30μM GSK1016790A n=2 of 3 tested cells).

We then used cryo-electron microscopy (cryo-EM) to determine the 3D structure of hTRPV4 in the detergent-solubilized state. The baculovirus-expressed hTRPV4 channel was stably reconstituted in buffer containing the detergent glyco-diosgenin (GDN) and purified to homogeneity. In order to obtain information about the 4α-PDD molecular mode-of-action, hTRPV4 was incubated with an approximate 10-fold molar excess of 4α-PDD and the structure of the resulting sample was solved using cryo-EM. Due to the limited aqueous solubility of 4α-PDD and the need to minimize DMSO effects during grid freezing, higher working concentrations of 4α-PDD were not achievable.

After optimization of the cryo-EM grid freezing conditions, hTRPV4 particles nicely entered thin ice and a good particle distribution in recorded images was obtained (**Extended Data Fig. 2a**). Two-dimensional class averages or extracted particle images showed that the sample adopted diverse orientations on the grid and secondary structural features were easily discernable (**Extended Data Fig. 2b**). Analysis and 3D sub-classification of the hTRPV4 particles indicated the presence of several conformational states (**Extended Data Fig. 2d**). In this contribution we present our analysis of the major conformational state which displayed features associated with a single tetrameric assembly. A four-fold rotational C4 symmetry was applied during the final stages of the cryo-EM structure reconstruction and the structure was refined at a global resolution of 4.1Å (**Extended Data Fig. 2e, Extended Data Table 1**). Despite significant differences in the local resolution throughout the channel (**Extended Data Fig. 2g**), map quality allowed unambiguous placement of all secondary structural elements and large bulky side-chains confirmed the correct registry assignment throughout the protein sequence. The resulting cryo-EM 3D reconstruction represents the first high-resolution hTRPV4 protein structural data (**Fig. 2a**).

**Extended Data Fig. 2.**
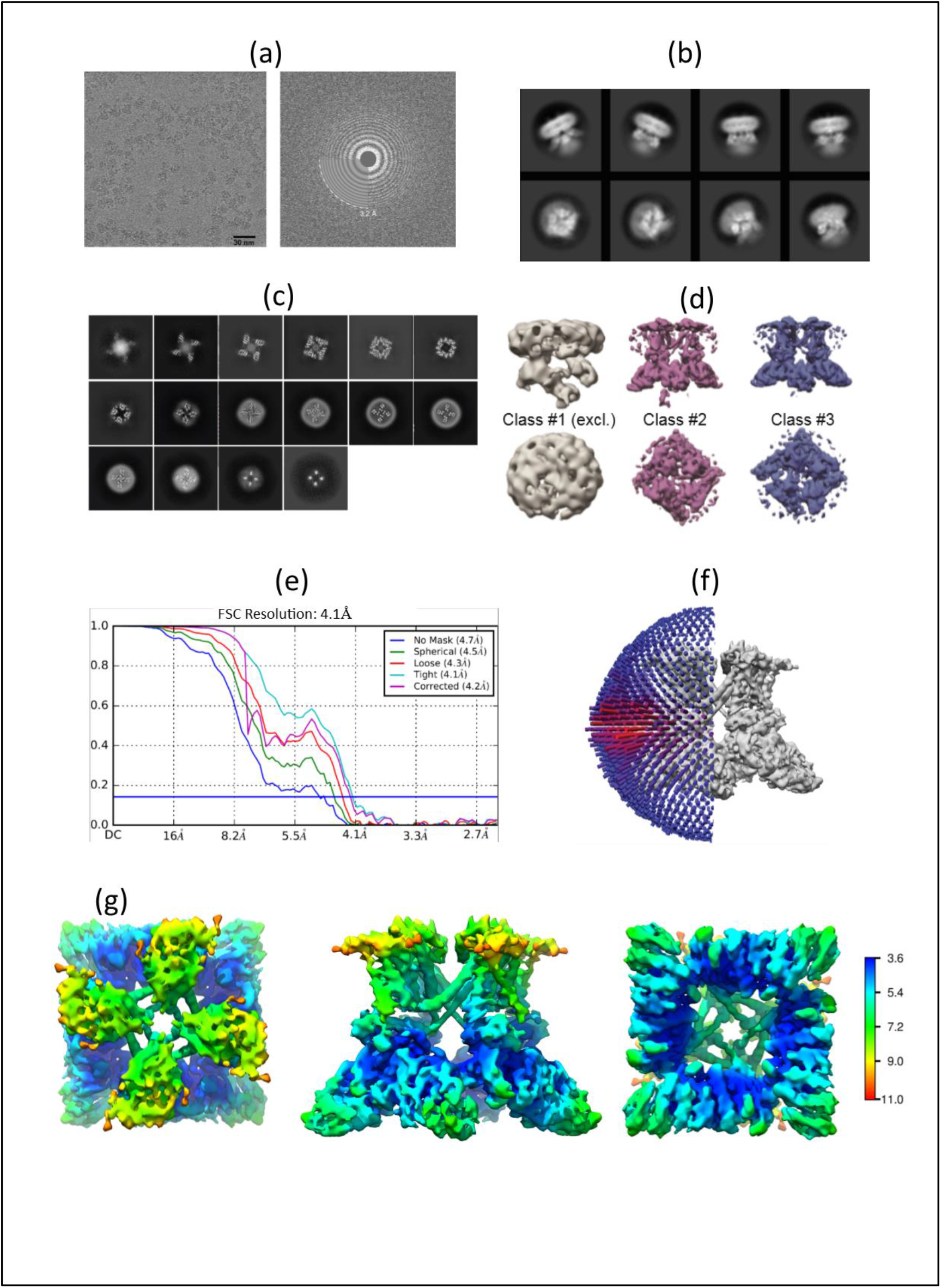
Single particle cryo-EM analysis of detergent solubilized hTRPV4 in presence of 4α-PDD. **(a)** Typical raw micrograph of detergent solubilized hTRPV4 in presence of 4α-PDD and corresponding computed power spectrum of the micrograph. (**b**) Representative 2D class averages. (**c**) Slabs of the unsharpened density map at different levels along the pore channel axis. (**d**) Classes obtained following 3D classification. Class 1 was poorly resolved. Classes 2 and 3 displayed similar features and were merged and subsequently refined. (**e**) Fourier shell correlation (FSC) curves after the final step of refinement. So-called gold standard protocols were used. The horizontal blue line indicates the applied 0.143 threshold for resolution estimation [38]. (**f**) Euler angle distribution for all the particles used in the final reconstruction. The position of each cylinder (blue) regarding the EM map (gray) indicates its angular assignment, whilst the cylinder height and color (blue to red) reflects the total number of particles in this specific orientation. (**g**) Cryo-EM map colored by local resolution, indicated in the scale in Å.

**Fig. 2.**
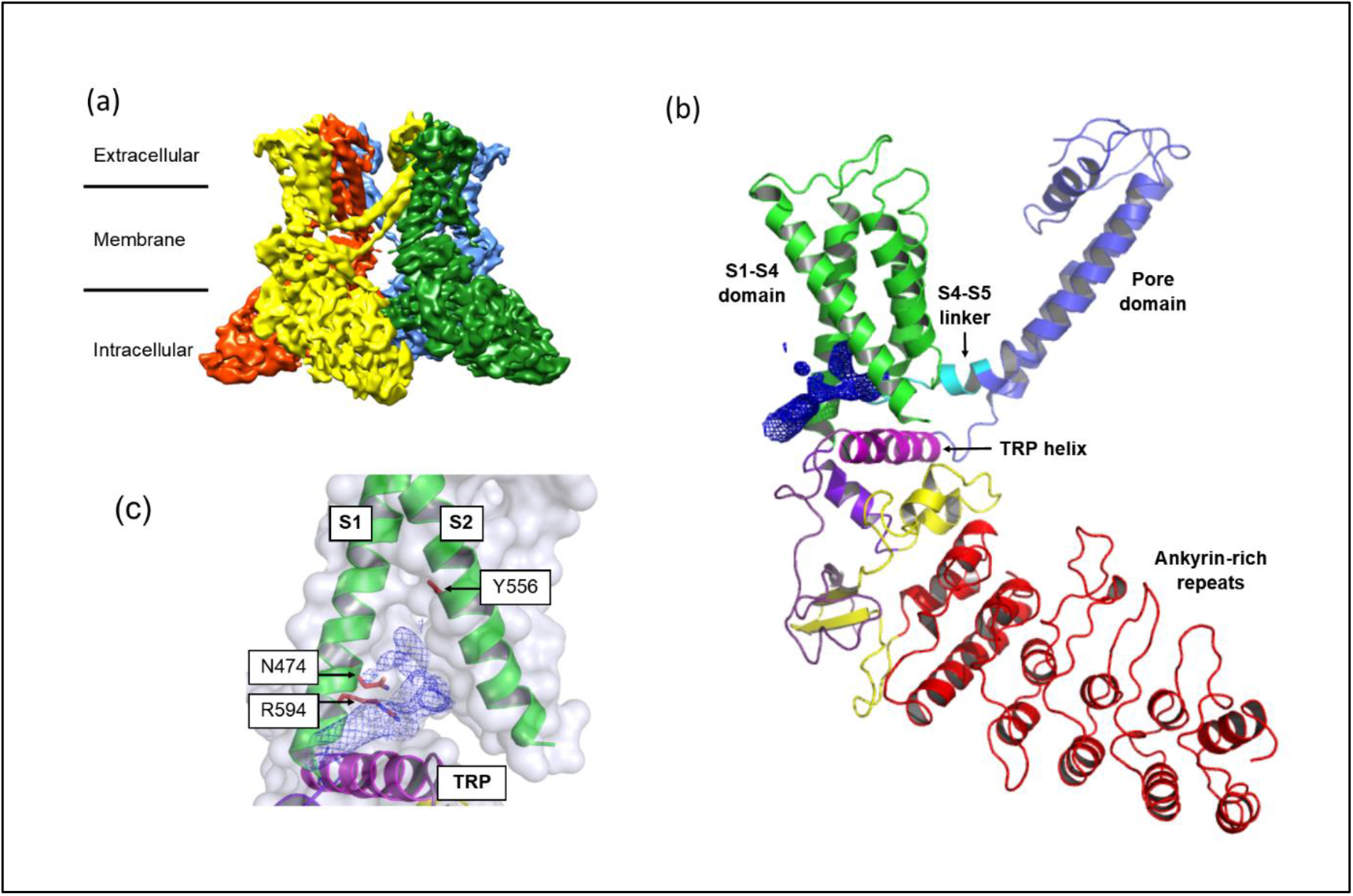
Cryo-EM structure of hTRPV4. (**a**) hTRPV4 cryo-EM reconstruction at 4.1 Å resolution with individual domain-swapped hTRPV4 subunits colored yellow, orange, blue and green displayed using the software UCSF Chimera [39]. (**b**) Monomeric view of hTRPV4 with individual structural elements labelled. (**c**) Observed non-protein cryo-EM map feature in the S1-S4 binding pocket (blue mesh).

### Overall architecture of hTRPV4

hTRPV4 adopts the archetypical TRP channel tetrameric assembly comprising a central transmembrane domain flanked by extra- and intra-cellular domains (**Fig. 2a**). The membrane-spanning TMD domain comprises six alpha-helices (S1-S6) and a TRP domain together with several adjacent helices in the elbow and pore region (**Fig. 2b**). The S1-S6 helices are ordered into two sub-domains whereby helices S1-S4 form an alpha-helical VSLD bundle that is flanked by the pore-lining S5-S6 helices in a domain-swapped arrangement. The adjacent intracellular domain comprises the ankyrin-repeat domain (ARD) and a helix-loop-helix region from the N-terminus of the protein, together with a helical-coiled domain from the C-terminus of the protein. Based on the over-stoichiometric presence of Ca^2+^ in the purification buffer, together with reports of cations binding at a similar position in other TRP channels, we modelled a single Ca^2+^ cation into the EM map in the upper selectivity-filter (SF) gate. The calcium ion is located within the SF-gate where it is flanked by carbonyl oxygen atoms from M681 that are located approx. 5 Å away. The presence of a single cation binding site is consistent with the classification of hTRPV4 as a non-selective cation channel. In contrast, cation selective ion-channels, for example TRPV6, obtains selectivity through multiple ion-binding sites within the ion channel pore [5]. The resulting hTRPV4 SF gate constriction point has a radius of approximately 10Å, a similar size to that observed in the xTRPV4 structure [25].

### 4α-PDD binding site

We hypothesized that an additional, strong non-protein feature in the cryo-EM map at the interface between the S1, S2 and TRP helices within the VSLD site represented the binding of 4α-PDD to hTRPV4 (**Fig. 2c**). The limited local resolution in this region of the map (**Extended Data Fig. 2g**), together with the inherently high degree of 4α-PDD structural flexibility, did not allow unambiguous modelling of the 4α-PDD binding mode. In order to validate this binding site, we therefore designed and characterized a series of hTRPV4 point mutants targeting both this and other known activity-modulating binding sites (**Table 1**). To allow direct experimental comparison with several previously reported 4α-PDD disrupting mutations, we also generated and tested Y556A, L584M, W586A and R594A hTRPV4 variants [35, 40]. Analysis of the open conformation hTRPV4 structure revealed that these residues are not all located within the same binding site, suggesting both direct and indirect effects on 4α-PDD activity.

**Table 1:** Impact of point mutations on 4α-PDD efficacy. EC_50_’s for structure-inspired point mutagenesis experiments as assessed by the impact on efficiency of 4α-PDD to open the channel in a fluorescent-based reporter assay. Maximal applied agonist concentration was 25 μM. Data are represented as mean ± standard deviation.

| Mutant | EC<sub>50</sub> (μM) |
| --- | --- |
| WT | 0.183 ± 0.092 |
| N474A | 1.04 ± 0.34 |
| N474Q | > 25 |
| K535A | 0.91 ± 0.26 |
| S548V | > 25 |
| F549A | 0.327 ± 0.081 |
| Q550A | 1.85 ± 0.66 |
| Y556A | 2.40 ± 0.53 |
| L584M | 0.67 ± 0.36 |
| W586A | 21.3 ± 6.4 |
| R594A | >25 |

All hTRPV4 point mutants were transiently expressed in CHO cells and their impact on activation via 4α-PDD was assessed in a calcium flux assay (Table 1, Extended Data Fig. 3). Dose-response curves were fitted to obtain EC_50_ values following 4α-PDD treatment. In initial experiments with full-length hTRPV4^WT^ we observed an EC_50_ of 0.18 μM upon treatment with 4α-PDD in good alignment with reported literature values [32, 35]. Similarly, literature reported trends for effects of the Y556A, L584M, W586A and R594A mutations were also observed in our calcium flux experiments [35].

From the tested panel of mutants, the strongest effects were observed for N474Q, R594A, S548V and W586A, all of which resulted in complete or almost complete loss of 4α-PDD induced Ca^2+^ influx. (**Table 1, Extended Data Fig. 3**). Furthermore, the N474A mutant also displayed an increased EC_50_ after treatment with 4α-PDD, albeit less strong than the N474Q mutant. Both N474 and R594 are located directly adjacent to the 4α-PDD density, supporting the identification of the 4α-PDD binding site (**Fig. 2c**). In contrast, W586 is located at the inter-subunit interface approximately 15 Å from the proposed 4α-PDD binding site, where it closely interacts with the S5 helix from an adjacent hTRPV4 protomer within the tetrameric assembly (**Extended Data Fig. 4**). Since W586 is surface exposed in the closed TRPV4 conformation, we hypothesize that the effects of the W586A mutation result from disruption to the formation of the open hTRPV4 conformation and not via direct disruption of 4α-PDD binding. It is, however, important to note that such conclusions cannot be fully supported by calcium flux measurements alone. More detailed electrophysiological characterization experiments will be required to fully characterize the disrupting mechanism of this mutation. The fourth mutant with strong disruptive effects, S548V, was discovered serendipitously. It is located within the S2/S3 loop adjacent to the putative 4α-PDD binding site. Whilst the loop is close to the proposed binding site, a detailed molecular rationale of its effect has not been possible, since this loop is not clearly resolved in the maps. A high degree of flexibility within the S2-S3 loop has also been observed in other TRP channel structures (**Extended Data Fig. 5)**.

The other tested mutants displayed, comparatively, only weak to modest effects on 4α-PDD-mediated channel activation. The Q550A mutation is located in the vicinity of the 4α-PDD binding site, and whilst the main-chain residues could be clearly modelled, the side-chain could not be unambiguously built. However, considering its location and likely side-chain conformations, it is conceivable that the Q550 mutation disrupts 4α-PDD activity. The Y556A mutation, located at the back of the 4α-PDD binding site directly at the interface between the S1 and S2 helices, resulted in an approximate 13-fold increase in the EC_50_. Two additional mutants were characterized to probe the impact of mutating the vanilloid binding or EET site on 4α-PDD activity. These mutants, F549A and K535A, resulted in either weak, or not significant changes to the 4α-PDD mediated activation and therefore suggest that 4α-PDD activity is not mediated via direct interactions with these alternative activity modulating sites. Collectively, these data support the identification of the 4α-PDD binding site at the interface between S1/S2/TMD helices within the VSLD-binding site.

**Extended Data Fig. 3.**
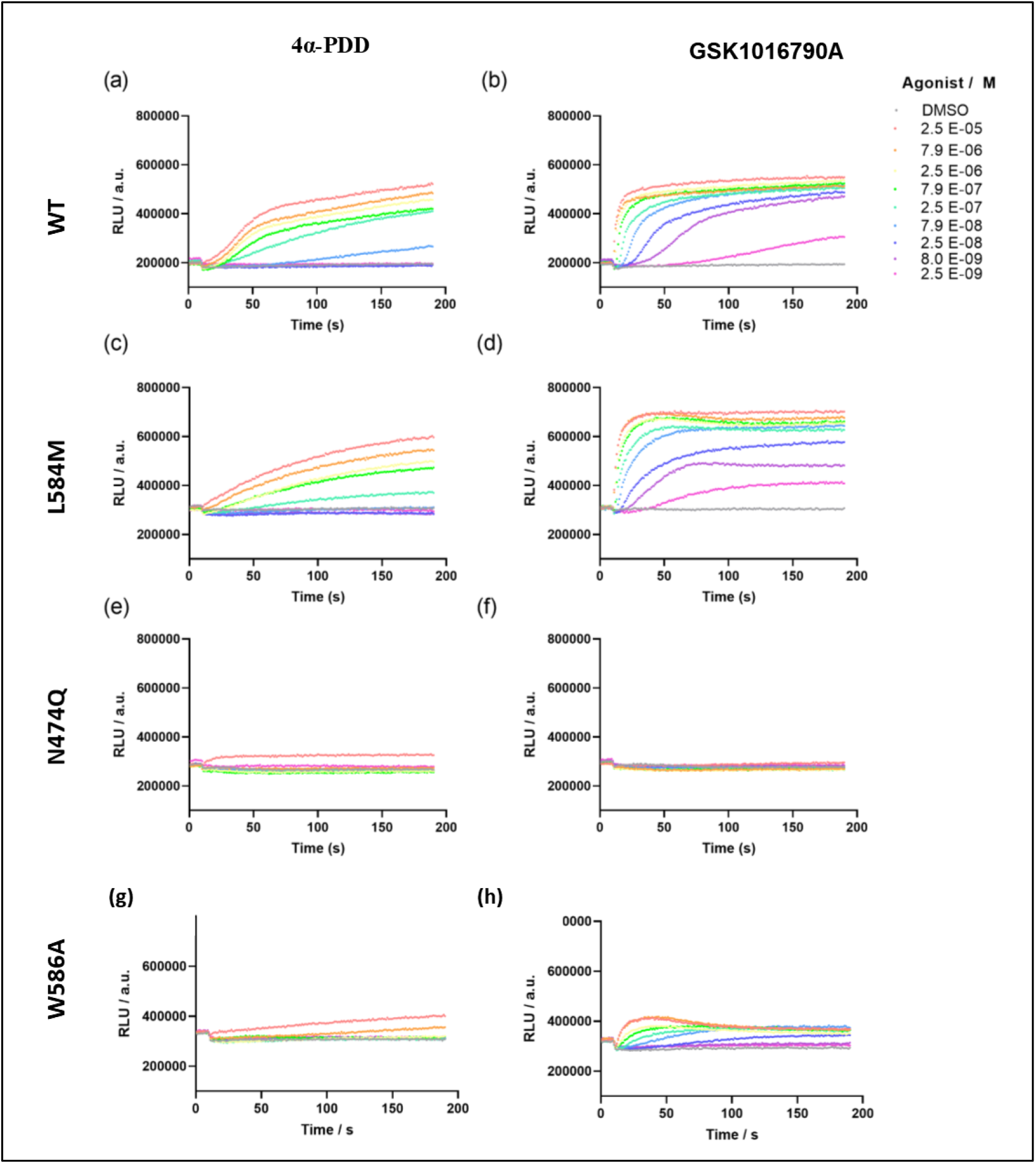
Exemplary hTRPV4 calcium flux data in response to 4α-PDD or GSK1016790A stimulation. Agonist is added 5s after the start of the measurement and Fluo8 fluorescence was recorded for 190 s as a measure of calcium flux. Highest concentration of agonist was 25 μM and agonists were diluted in half-log dilution steps. A-B show the response of the hTRPV4^WT^ towards the agonists. C-D show the response of the L583M mutant in the vanilloid site leading to little modulation compared to the wt. E-F show the mutant N474Q causing a strong effect abolishing the calcium response of both agonists. G-H show the W586A mutant causing a strong decrease in potency and a strong decrease in calcium influx.

**Extended Data Fig. 4.**
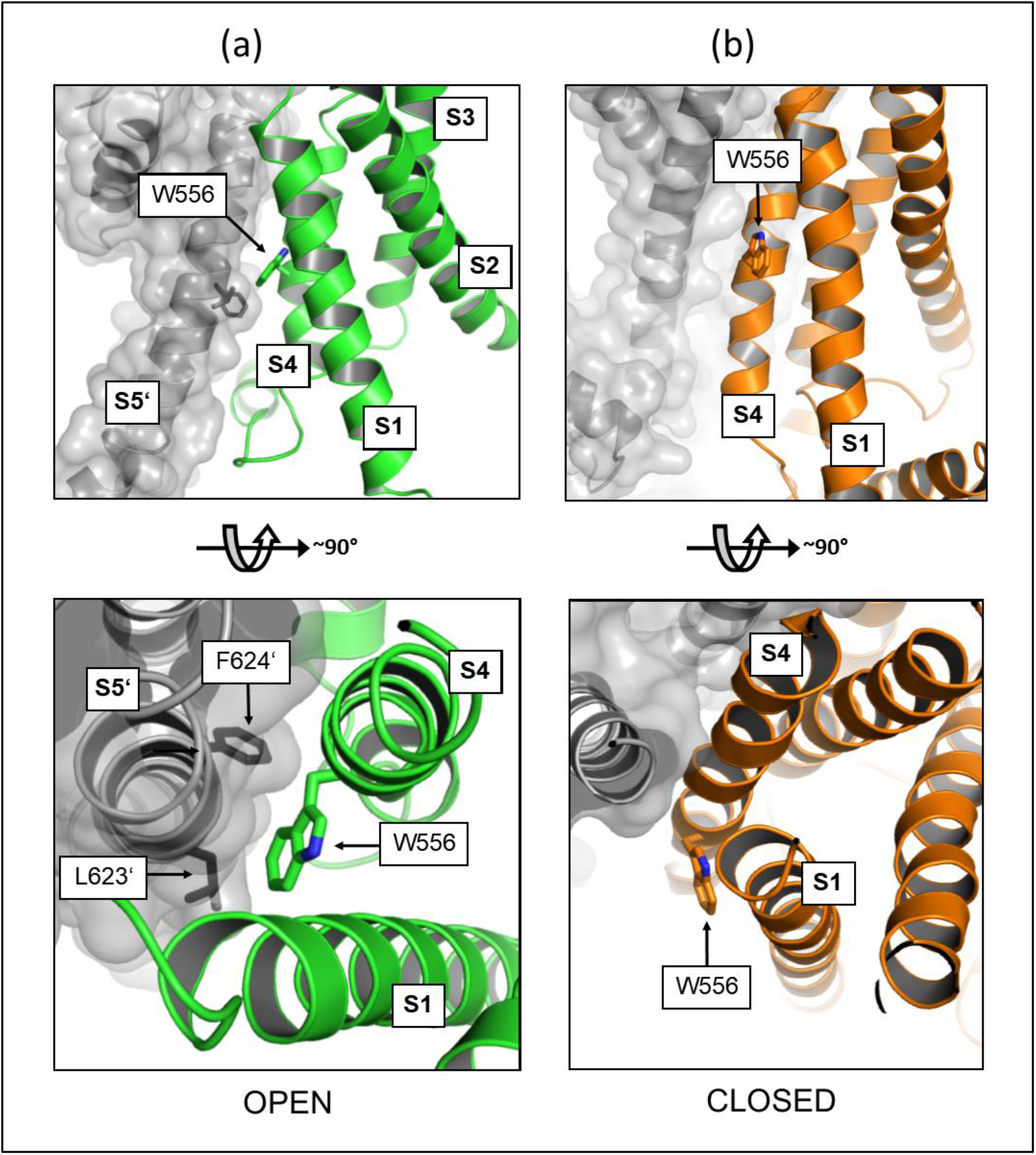
Structural role of hTRPV4 W586. (**a**) Orthogonal views of the W596 side chain location in the open-conformation 4α-PDD bound structure. The S1-S4 bundle of one protomer is shown is depicted in a green cartoon representation. The neighboring, strand-exchanged protomer, is shown in gray. (**b**) Orthogonal views of the W596 side chain location in the closed-conformation 4α-PDD bound structure. The S1-S4 bundle of one protomer is shown is depicted in an orange cartoon representation. The neighboring, strand-exchanged protomer, is shown in gray.

We also characterized the impact of these mutants on GSK1016790A mediated hTRPV4 activation (Extended Data Table 2, Extended Data Fig. 3). As for 4α-PDD, the observed TRPV4^WT^ EC_50_ of 1.2 nM is in good alignment with the literature-reported value of 2.1 nM [33]. Similar to 4α-PDD, N474Q and R594A mutations completely abolished any calcium response to GSK1016790A. The W586A mutation also significantly affected the GSK1016790A response leading to a reduced calcium flux indicated by the lower fluorescence increase. In contrast to the 4α-PDD, however, it was still possible to resolve an EC_50_ of 0.37 μM which is factor ~20 increased compared to the wildtype channel. The weak calcium influx indicates that, in contrast to the weak 4α-PDD agonist, this mutation does not completely disrupt the open conformation upon stimulation by the stronger GSK1016790A agonist. Taken together, the reduced potency of the agonists and the reduced calcium flux may indicate changes to the open conformation. Interestingly, the impact of the W596A mutation has also been observed to differ between other TRPV4 stimuli. For example, whilst W586A also disrupts the channel sensitivity to bisandrographolide A and heat, the mutation has no effect on the channel response to cell swelling, arachidonic acid and 5,6-EET [35]. As for 4α-PDD, vanilloid-binding sites mutants displayed only weak effects on GSK1016790A potency, changing the potency by less than an order of magnitude. Taken together, these shared trends suggest an overlap between the 4α-PDD and GSK1016790A binding sites. Future in-depth electrophysiological characterizations, which are outside the scope of this project, will be required to disentangle the molecular mechanisms.

**Extended Data Table 2.** Impact of point mutations on GSK1016790A efficacy. EC_50_’s for structure-inspired point mutagenesis experiments as assessed by the impact on efficiency of GSK1016790A to open the channel in a fluorescent-based calcium flux assay. Maximal applied agonist concentration was 25 μM. Data are represented as mean ± standard deviation.

| Mutant | EC<sub>50</sub> (nM) |
| --- | --- |
| WT | 1.210 ± 0.033 |
| N474A | 540 ± 150 |
| N474Q | >25000 |
| K535A | 7.4 ± 2.9 |
| S548V | 110 ± 18 |
| F549A | 9.8 ± 4.1 |
| Q550A | 27 ± 13 |
| Y556A | 9.6 ± 4.3 |
| L584M | 4.0 ± 1.7 |
| W586A | 36 ± 21 |
| R594A | >25000 |

### Structural changes upon hTRPV4 activation

To investigate the molecular basis of 4α-PDD induced activation we next built an apo, closed conformation hTRPV4 homology model using a closed conformation xTRPV4 structure as a template [25]. Comparison of the pore structure of this model with our 4α-PDD-complexed hTRPV4 structure revealed that whilst the upper gate architecture is retained between the two states, the lower gate region changes significantly. Transition to the open-state is accompanied by an approximate 90° counter-clockwise rotation of the S6 helix (as viewed from the extracellular side), resulting in I715 replacing M718 as the residue defining the constriction point (Fig. 3a/b, Suppl. Movie S1). The pore diameter at the lower gate increases considerably from 5.4 Å in the closed state (M718 Cα residues) to 10.6 Å in the open state. A translation of the S6 helix, together with the smaller size of I715 compared to M718, translates into an increased van der Waals radius (from 0.8 Å to 1.8Å) and thus movement of a hydrated ion through the channel is possible (Fig. 3c). We therefore conclude that in the presence of 4α-PDD, hTRPV4 is stabilized in an open ion-channel conformation.

**Fig. 3.**
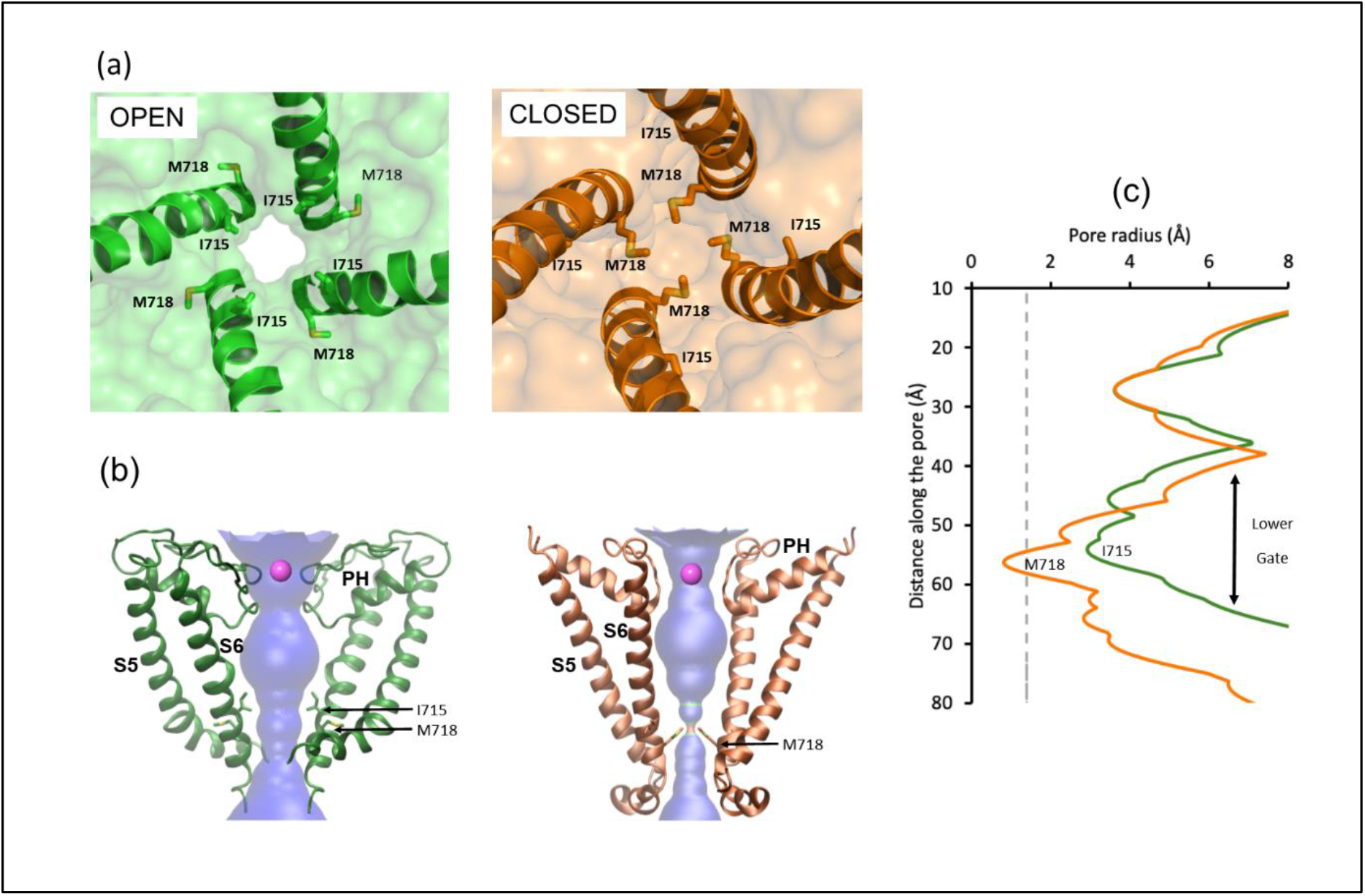
Open and closed state hTRPV4 ion pore structures. (**a**) Cartoon representation of the open-state lower gate constriction point (green) and the equivalent position in the hTRPV4 closed state model (orange) as viewed from the extracellular side. The S6 helix is depicted in cartoon representation and key residues are highlighted in stick representation, with C atoms colored as in the parent structure and N and O atoms colored blue and red, respectively). (**b**) Cartoon representation of the open (green) and closed (orange) pore profile generated using the HOLE software [41]. A bound Ca^2+^ ion in the upper selectivity filter region (pink) and other key residues are highlighted as described in (a). (**c**) Graphical representation of the radius of the 4α-PDD bound open (green) and closed (orange) state hTRPV4 pore profile. The dotted line represents the van-der-Waals radius of a water molecule (1.4 Å). Specific residues lining the pore constriction points are labelled.

The altered pore properties result from large conformational changes to the TMD region such that the relative orientation of the S1-S4 bundle and the pore-forming region are strikingly different in the open and closed states. Compared to the closed state, the open state S1-S4 helical bundle is rotated approximately 90° (clockwise, as observed from extracellular side) about the S4 helical axis (**Fig. 4a, Suppl. Movie S2**). The internal arrangement of helices within the S1-S4 domain itself does not change between open and closed conformations (rmsd of 1.7Å for residues 467-594). In contrast, interactions between the S1-S4 helical bundle and several flanking structural channel elements change significantly. Most strikingly, upon channel activation the TRP helix orientation changes by approximately 60°, such that it then forms an integral part of the 4α-PDD binding pocket. The TRP helix is positioned directly between the S1 N- and S2 C-termini, orthogonal to the plane of the two helices, and is therefore ideally positioned to form direct interactions with 4α-PDD in the open conformation (**Fig. 4b, Suppl. Movie S3**). In contrast, the TRP helix is located further away from the S1/S2 helices in the closed state and does not contribute directly towards the 4α-PDD binding site. Due to the altered TRP helix conformation, the S2-S3 linker cannot adopt its closed-state conformation. A high amount of structural flexibility in this region prevented us from unambiguously building this linker. Additionally, the unique TRPV4-closed conformation tight hydrophobic interface between S3/S4 and S5/S6 is disrupted in the open-conformation. The reduced intimacy between the TMD and pore-forming domains is in turn associated with pronounced changes to the open state pore-forming region. A kink in S5 is observed (around K612), such that the N-terminal region of the helix, at around the same height as the lower gate, is positioned further away from the pore. This facilitates a widening of the lower gate through an altered S6 conformation.

**Fig. 4.**
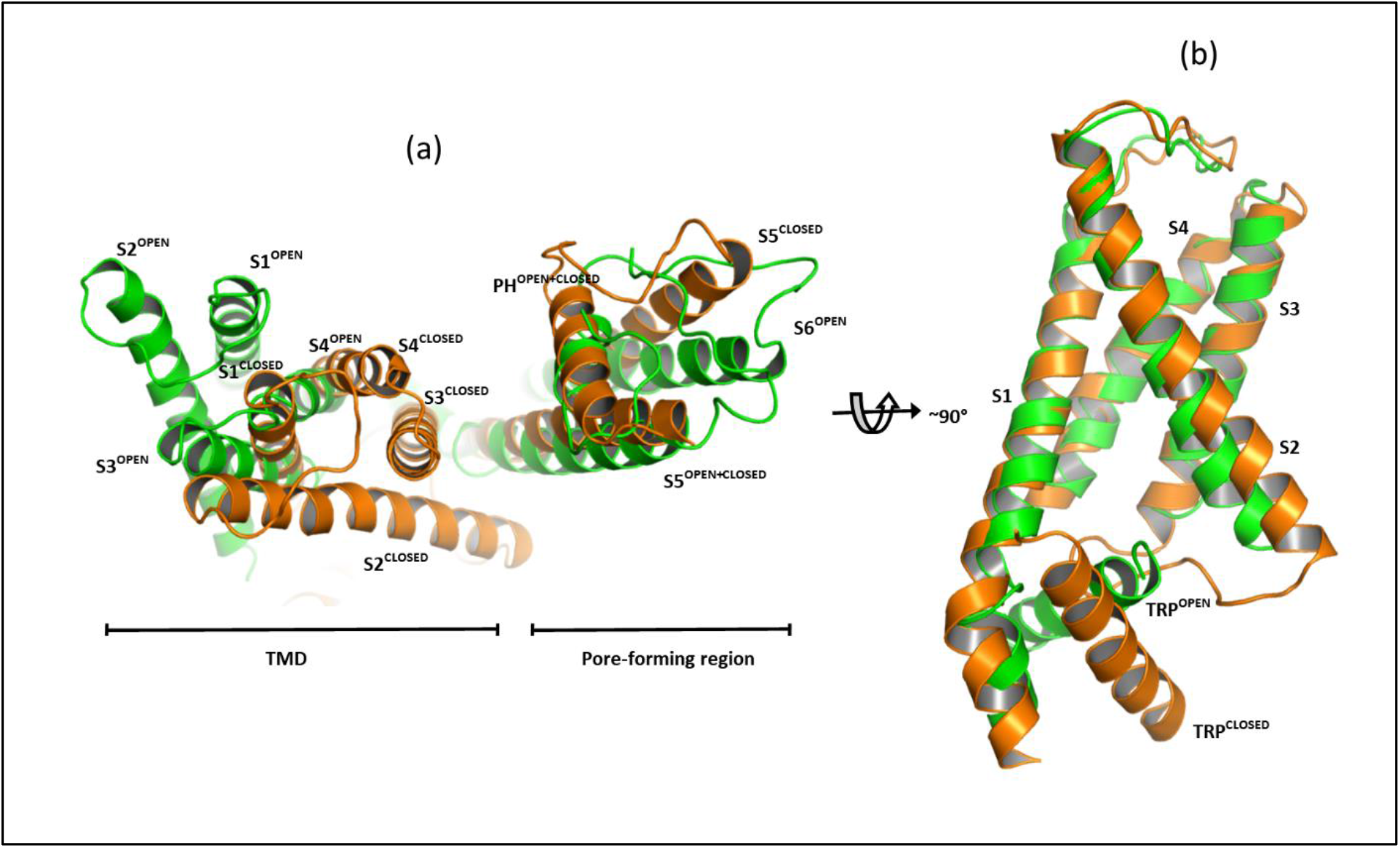
hTRPV4 conformational changes upon 4α-PDD binding. (**a**) Overlay of the hTRPV4 4α-PDD (green) and hTRPV4 closed model (orange) aligned based on the tetrameric pore (S5-S6) helices. (**b**) Overlay of TMD domains from hTRPV4 4α-PDD (green) and the hTRPV4 closed model (orange) aligned based on the S1-S4 helical bundle.

Collectively, the binding of 4α-PDD appears to trigger structural rearrangements in the TRP-helix orientation that then propagate though the protein and ultimately result in channel opening. Indeed, the TRP helix has previously been described as a structural element – or force hub - that integrates allosteric signals from different channel domains into the pore [42]. In particular, the interaction between the TRP helix, S4-S5 linker and S6 helix elements and the transmission of stimuli to the gate has been well-characterized [28]. The resulting overall open-conformation of hTRPV4 generally resembles that of other open -state thermo-TRP channels (Extended Data Fig. 5), with the TMD domains of active TRPV1-3 structures sharing RMSDs in the range of 2-3 Å. Thus, whilst TRPV4 displays a non-archetypal closed, or inactive conformation, its active conformation extends the TRP channel members for which the active conformation TRP channel paradigm is conserved (Extended Data Fig. 6).

**Extended Data Fig. 5.**
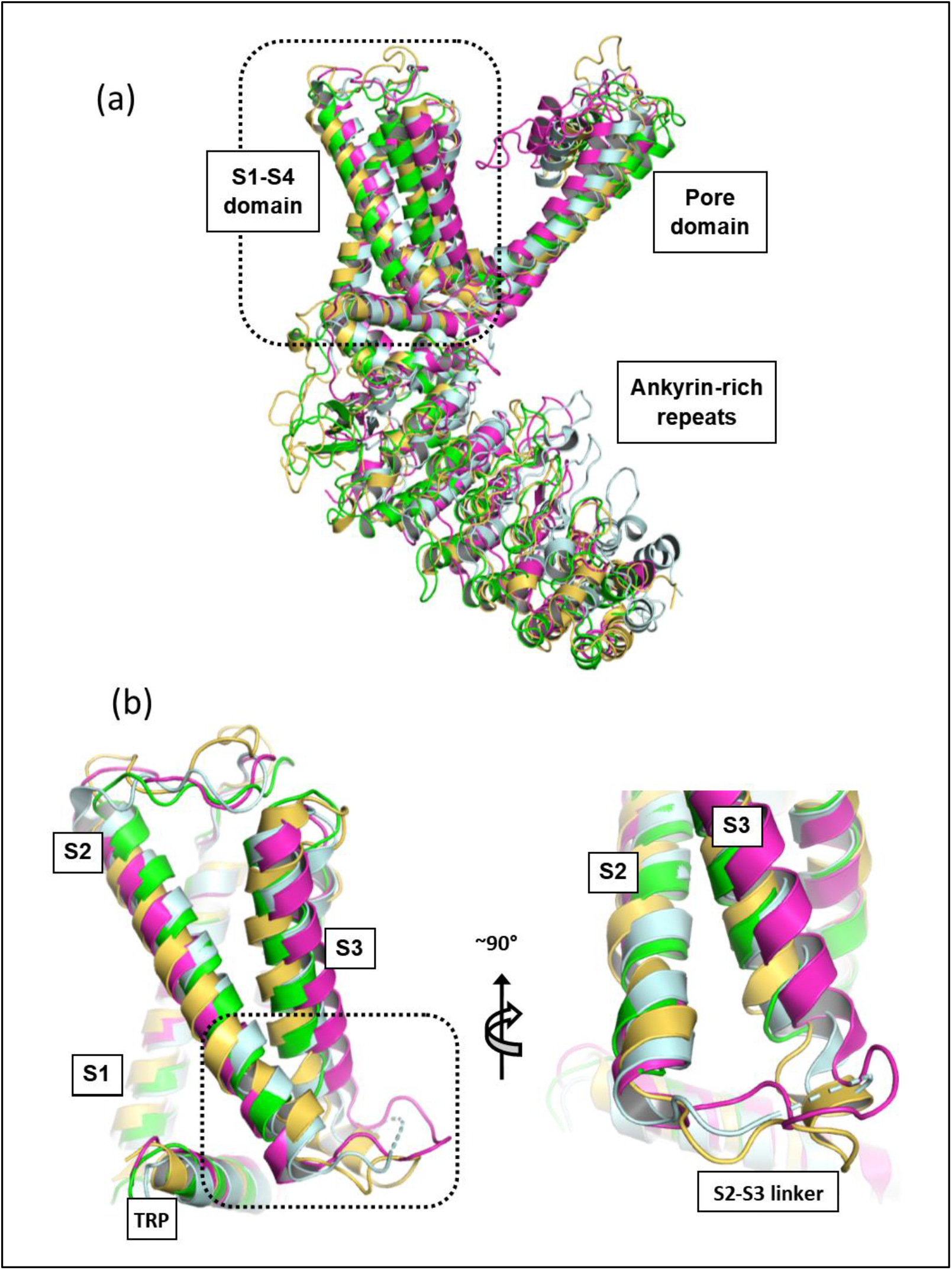
Superposition of hTRPV4 TMD with other open conformation thermo TRP channel structures. (**a**) Overlay of TRPV1 (PDB code 3J59, light blue), TRPV2 (6BOV, magenta), TRPV3 (6DVZ, yellow) and hTRPV4 (this publication) open conformation structures based on residues within the TMD domain. (**b**) Orthogonal views showing a close-up of the S2-S3 linker region, colored as in (a).

**Extended Data Fig. 6.**
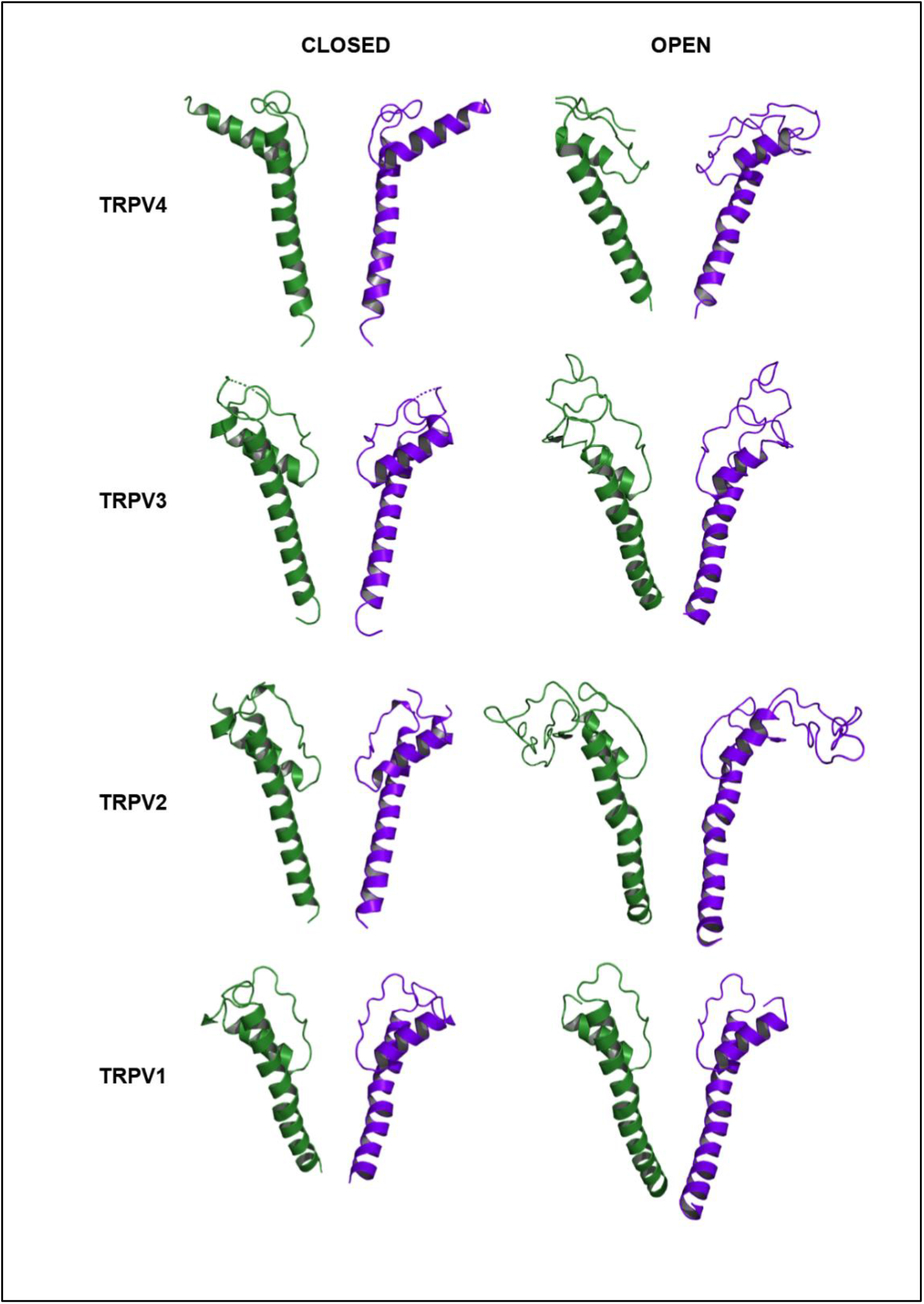
Open and closed conformation of the pore in TRPV1-4 representatives from different organisms. The pore helix and S6 helix are shown from 2 different monomers (shades of green and blue). Closed conformation structures (left-hand side): xTRPV4 (PDB code 6BBJ), human TRPV3 (6MHO), rabbit TRPV2 (5AN8) and rat TRPV1 (3J5P). Open conformation structures (right-hand side): hTRPV4 (this study, 7AA5), mouse TRPV3 (6DVZ), rat TRPV2 (6BO4) and rat TRPV1 (3J5Q).

### 4α-PDD binding-mode model

To gain further insights into the 4α-PDD molecular mode of action, we next performed 4α-PDD docking calculations guided by the EM map. Importantly, these calculations confirmed that the binding site is large enough to accommodate the large 4α-PDD molecule. Due to the inherent high degree of structural flexibility within the 4α-PDD molecule, a large number of candidate poses within the hTRPV4 binding pocket were identified during the sampling phase of the docking algorithm that were narrowed down to five by applying restraints to improve the fit between the proposed binding mode and the EM map. The top five scoring poses displayed a similar placement of the 4α-PDD phorbol group and differed only by the detailed positioning of the acyl chains escaping from the binding site toward the membrane. Based on its favorable docking score, a putative binding mode was identified that is consistent with both our mutagenesis data and published 4α-PDD structure-activity relationships [32] (**Fig. 5**). In this model, the core diterpenoid moiety binds at the interface between the S1, S2 and TRP helices, where it forms specific hydrogen bonds with protein residues within the binding site, most notably with N474 (**Extended Data Fig. 7a**). The long and highly flexible lipophilic alkyl chains extend out of the pocket into the areas embedded within the hydrophobic membrane environment. This is consistent with published SAR studies indicating that the acyl chains are involved in positioning the diterpenoid core for binding, rather than interacting with the binding site [32]. Furthermore, the sequence conservation between the different thermo TRPs within this pocket shows divergent residues lining the site (**Extended Data Fig. 7b)**, consistent with 4α-PPD activity with TRPV4, but not TRPV2 and TRPV3 [11].

**Fig. 5.**
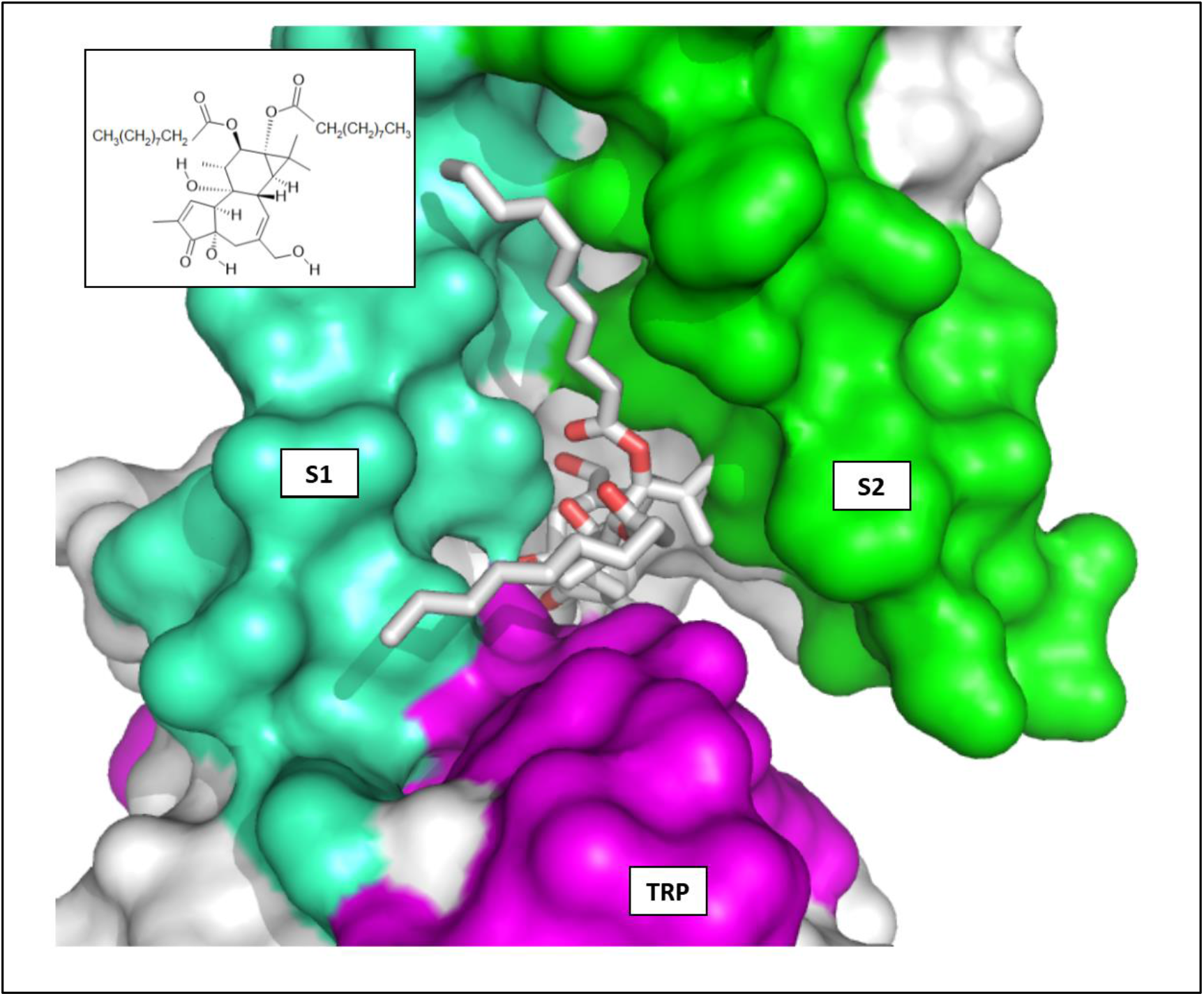
4α-PDD binding to hTRPV4. Cryo-EM map guided docking model of 4α-PDD binding in the hTRPV4 VSLD site. Surface representation of hTRPV4 with the S1, S2 and TRP helical elements colored green-cyan, green and magenta, respectively. 4α-PDD is shown in stick representation with C and O atoms colored white and red, respectively. The inlet depicts the structure formula of 4α-PDD.

**Extended Data Fig. 7.**
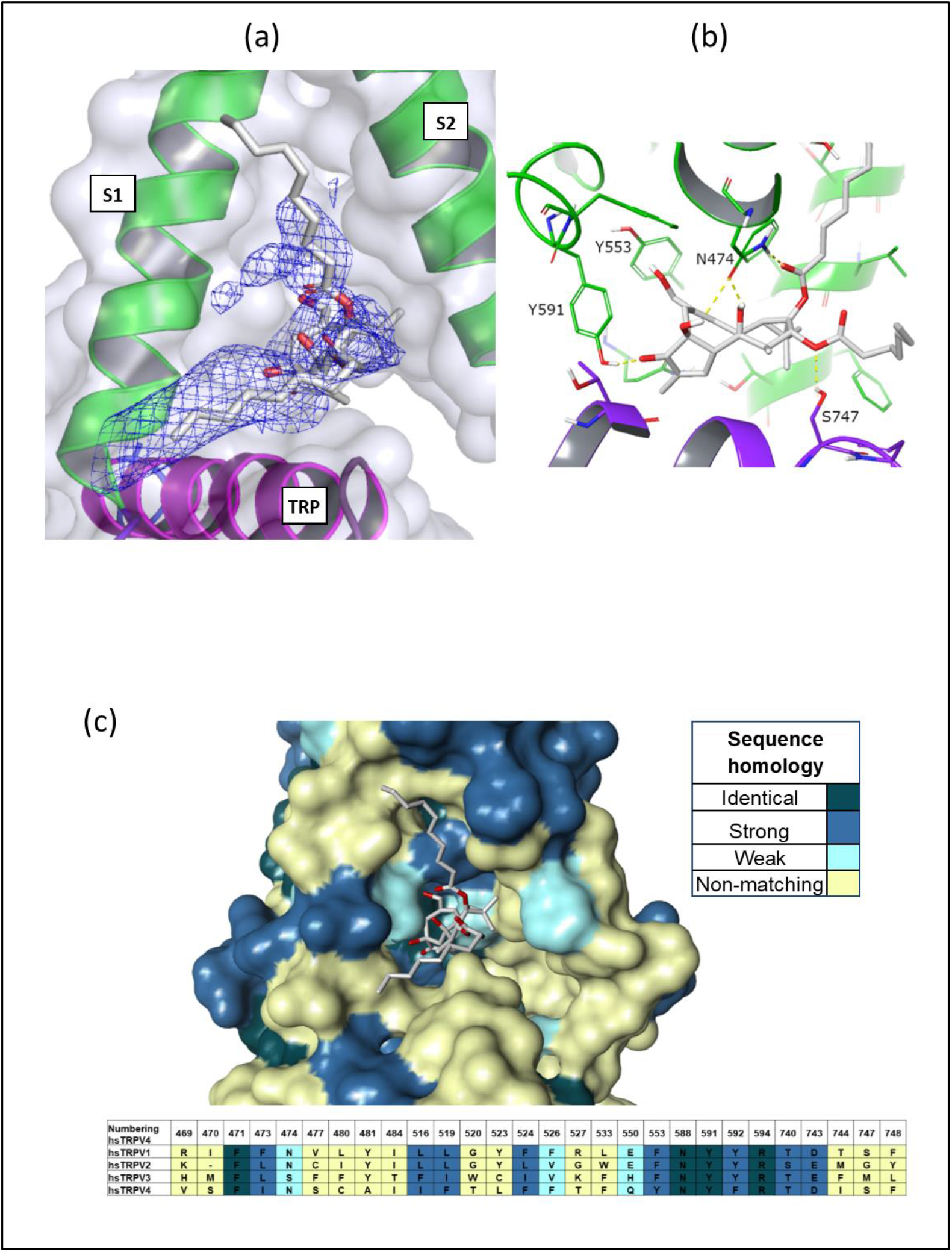
4α-PDD binding model. (**a**) Docking of 4α-PDD into the non-protein difference cryo-EM map feature in the VSLD binding site. The 4α-PDD molecule carbon and oxygen atoms are colored white and red, respectively. (**b**) Molecular details of the interaction between hTRPV4 and the docked 4α-PDD molecule, colored as in (a). (**c**) Sequence homology between hTRPV1, 2, 3, 4 sequences mapped onto the 4α-PDD complexed hTRPV4 structure. Residue identities and conservation for residues within 5Å of the binding site are shown in the alignment.

## DISCUSSION

The high-resolution cryo-EM structure of human TRPV4 in complex with 4α-PDD provides first insights into the agonistic molecular mode-of-action. Despite only moderate activity of the agonist in both, cellular electrophysiology and FRET assays, as well as in biophysical assays measured in detergent solubilized protein, the concerted mode-of-action of the four binding sites within the homo-tetrameric assembly results in large conformational changes to the ion-channel architecture that collectively result in channel pore opening. A combination of complementary structural and biochemical data allowed location of the allosteric 4α-PDD binding site within the S1-S4 bundle. This allosteric agonist binding site is perfectly located to facilitate the structural change from the closed to the open conformation through interactions with the TRP helix. Several structurally diverse small molecule effectors have now been observed to bind at this allosteric binding site in several members of the TRP-family of ion-channels (**Fig. 6**). Intriguingly, whilst compound binding at this site in the thermo-TRP family elicits agonistic effects, binding to the related TRPV5 and TRPV6 proteins results in antagonist effects. Obviously, small differences in the chemical structure of the small molecule at the binding site modulate very different overall structural response that leads to activation or inhibition of the channel. This concept is well known in the area of GPCR research and observed in CCK2, CCR2, 5HT, D2, opioid receptors and many more. For example, replacing the N-methyl group on potent m-opioid agonist morphine (Ki=0.53nM) with N-allyl resulted in a potent antagonist nalorphine (Ki=0.36nM). Furthermore, addition of an aliphatic side chain to a Ghrelin receptor modulator changed it from a 36nM agonist into a 20nM antagonist [43]. Consequently, the ligand binding site exploited by 4α-PDD in hTRPV4 exemplifies this activity switch for an ion-channel and more examples might be identified in the future due to the increasing structural information for this class of drug targets. Further high-resolution studies with potent agonists and antagonists at this binding site within hTRPV4 are required to further understand the mechanism of activity response and structural requirements.

The structural and biochemical data presented here are a valuable tool to support the further exploration of TRPV4 biology and its pharmacological responses, as well as the rational design of effective drug molecules. The interpretation of further functional characterization within the context of a three-dimensional target understanding promises to provide detailed insights into the processes regulating TRPV4 activity. Structural data has now identified several activity-modulating sites common to various TRP ion channel family members. Overall, this knowledge provides a great opportunity to support structure-guided design towards potent, selective and optimized novel drug molecules in a broad spectrum of human diseases.

**Figure 6.**
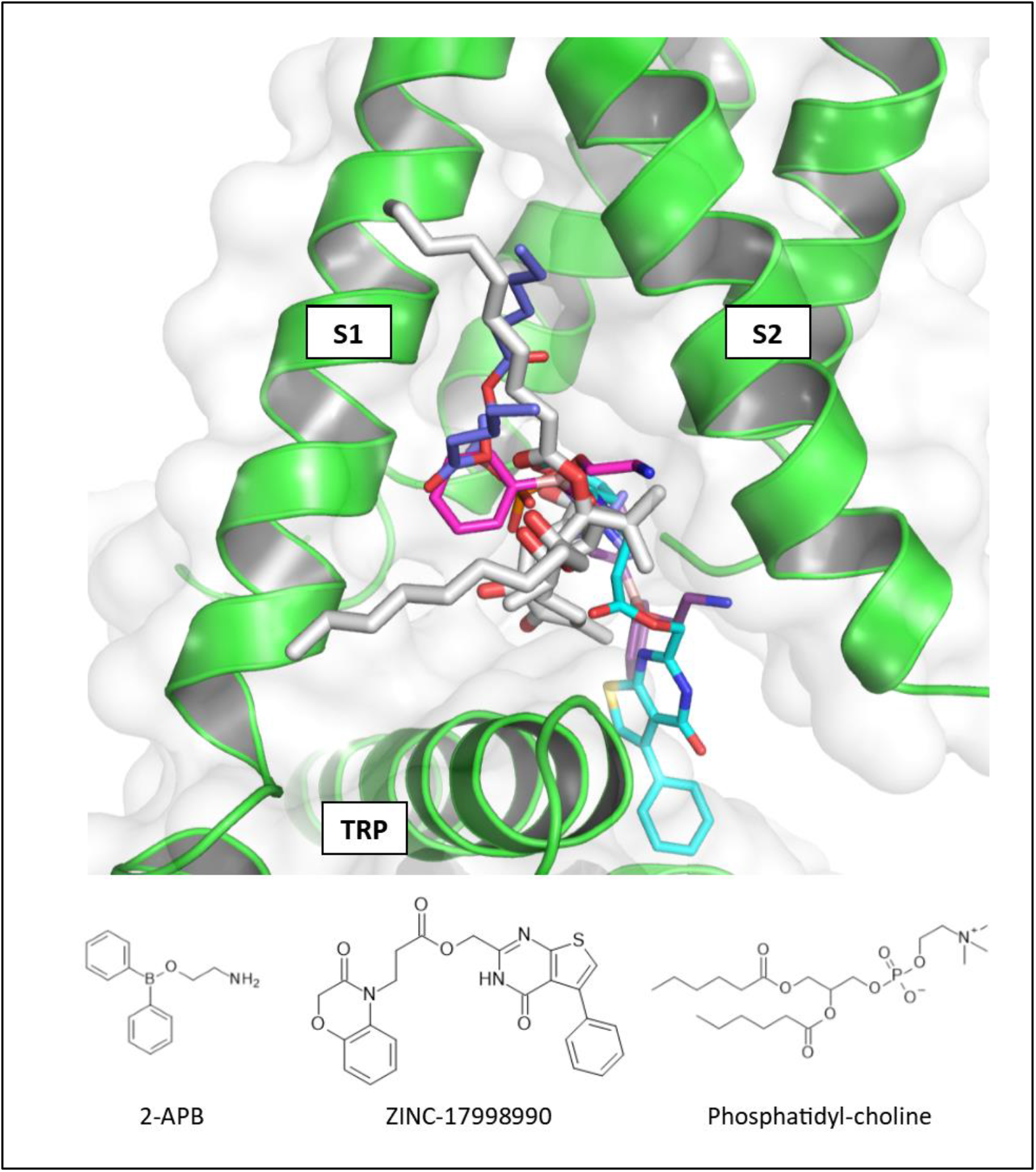
Ligands bound in the hydrophobic cleft between TRP channel S1 and S2 helices. Overlay of ligands binding to the TRP VSLD binding site based on the S1-S4 helical bundle. The following ligands are depicted: 4α-PDD/hTRPV4 (white), phosphatidyl-choline/TRPV1 (PDB code 5IRX, blue), Zinc-17998990/TRPV5 (6PBE, cyan), 2-APB/TRPV6 (6D7O, purple) and 2-APB/TRPV3 (6DVY, magenta). For clarity only the hTRPV4 protein binding site is depicted.

## Movie Legends

**Supplementary Movie S1 |** A movie from the extracellular side of the channel showing the transformation in the S6 helix conformation upon transition from the closed hTRPV4 conformation (orange) to the 4α-PDD bound open conformation (green) and back to the closed conformation (orange). Transition from the closed to open state is accompanied by a rotation of the S6 helix, such that I715 replaces M718 as the constriction point defining residue.

**Supplementary Movie S2 |** A movie showing the conformational changes to the TMD and pore-forming region upon transition from the closed hTRPV4 conformation (orange) to the 4α-PDD bound open conformation (green) and back to the closed conformation (orange). The structures are superposed based on the tetrameric pore (S5-S6) helices.

**Supplementary Movie S3 |** A movie showing the conformational changes to the TRP helical element upon transition from the closed hTRPV4 conformation (orange) to the 4α-PDD bound open conformation (green) and back to the closed conformation (orange). The structures are superposed based on the S1-S4 helical bundle.

## ONLINE METHODS

### Protein expression and purification

A synthetic hTRPV4 gene fragment encoding residues 148-787^N651D^ (predicted glycosylation site mutated) was integrated into the baculovirus pVL1393 transfer vector. The resulting construct additionally encodes an N-terminal FLAG tag followed by a C3 protease cleavage site and a C-terminal TEV cleavage site followed by eGFP and a terminal His10 tag. High-titer recombinant baculoviruses were obtained with the FlashBAC according the manufactures protocol. For large scale recombinant protein production Sf9 cells at densities of 4.0 x 10^6^ cells/mL were infected with high-titer viral stock at a multiplicity of infection (m.o.i) of 1.5. Cells were incubated for 48 h at 27 °C in a cell wave bag at 22 rpm. Cells were harvested by centrifugation, flash frozen and stored at −80 °C until use.

The overall purification steps were carried out at 4°C or on ice. Biomass corresponding to 4 liters of insect cell culture was thawed on ice and resuspended in lysis buffer (25 mM Tris pH 8.0, 150 mM NaCl, 2 mM CaCl_2_) supplemented with cOmplete™, EDTA-free protease inhibitor Cocktail (Roche Applied Science). Cells were lysed and homogenized by 4 passages through a LM10 Microfluidizer^®^ (Microfluidics™) at an operational pressure of 8,000 p.s.i. Membranes were isolated by centrifugation (150,000*g*, 45 min) and resuspended in solubilization buffer (25 mM Tris pH 8.0, 150 mM NaCl, 2 mM CaCl_2_, 20mM imidazole pH 8.0) supplemented with cOmplete™, EDTA-free protease cocktail inhibitor. Subsequent solubilization of membrane proteins was performed by addition of 1% (w/v) glyco-diosgenin (GDN, Anatrace) and incubation under gentle agitation at 4°C for 2 hours. The insoluble fraction was removed by centrifugation (150,000*g*, 45 min). Detergent solubilized hTRPV4 was captured on a 5 ml Ni-NTA Superflow Cartridge (Qiagen) pre-equilibrated with solubilization buffer supplemented with 1% GDN (w/v). Ni-NTA resin was washed with isolation buffer (25 mM Tris pH 8.0, 150 mM NaCl, 2 mM CaCl_2_, 0.01% (w/v) GDN) supplemented with 20 mM imidazole pH 8.0 followed by a further wash with isolation buffer supplemented with 40 mM imidazole pH 8.0. Protein elution was performed with isolation buffer supplemented with 300 mM imidazole. Immediately following elution, a desalting step was performed by passing the eluted pool of hTRPV4 over a PD-10 desalting column containing Sephadex G-25 resin (GE Healthcare) pre-equilibrated with isolation buffer. hTRPV4 was subsequently concentrated with a 100-kDa Vivaspin Turbo concentrator (Sartorius) to a final volume of 500 μl. The sample was subjected to a final size-exclusion chromatography using a Superose 6 Increase 10/300 GL column (GE Healthcare) pre-equilibrated with isolation buffer. Eluted peak fractions were quantified and typically contained 0.75-1.00 mg/ml of hTRPV4 homotetramer.

### Cryo-EM grid preparation and data acquisition

The peak hTRPV4 fraction was incubated with 4α-PDD at a final concentration of 100 μM for 30 minutes on ice. 3 μL of purified concentrated hTRPV4 4α-PDD complex was then applied to QF 1.2/1.3 grids (Quantifoil, Jena, Germany) holey carbon grids glow discharged in air for 30s. Grids were blotted for 2s and vitrified in liquid ethane using a Vitrobot mark IV (FEI company) operated at 8 °C and 90% relative humidity. 19114 movies were collected over 3 sessions on a FEI Titan Krios (ThermoFisher Scientific) operated at 300 kV and equipped with a Quantum-LS energy filter (slit width 20 eV) with a K2 Summit direct electron detector (Gatan Inc.). Setup of automated data collection was done in SerialEM [44]. Movies were recorded in electron-counting mode fractionating 50 electrons per square Angstrom over 40 frames and with a pixel size of 0.639 Å/px.

### Cryo-EM image processing and analysis

After initial dataset pruning with the software FOCUS [45], Cryosparc2 was used for subsequent processing [46]. Drift correction and contrast transfer function estimation were performed using alignparts_lmbfsg and CTFfind4.1 within Cryosparc2 [47]. Aligned averages with poor CTF estimation statistics or high drift profiles were discarded. Automated particle picking from the remaining 18,000 images resulted in 1,040,228 particle locations. Particles were extracted, Fourier-cropped to 4 Å/px and 2D classified. After several classification rounds, the best 301,000 particles were submitted to 3D classification by means of multi class *ab-initio* reconstruction and heterogeneous refinement. 134,854 particles belonging to the best resolved class were corrected for local motion and re-extracted. Further 3D classification in C1 symmetry resulted in two similar classes with clear secondary structure elements and 4-fold symmetry features and one poorly resolved class. 62,385 particles from the 2 well resolved classes were used in non-uniform refinement imposing C4 symmetry. The resulting map has an estimated resolution of 4.18 Å as judged by the FSC cutoff of 0.143 [38].

### Model building and refinement

Model refined was performed with the Phenix version 1.16-3549-000 real space refinement protocol, applying a 4-fold internal molecular symmetry using non-crystallographic symmetry (NCS) constraints together with secondary structure and Ramachandran plot restraints. Morphing and atomic displacement parameters (ADP) refinement were included as per program default setting. Except for two loop regions (residues 533-548 and 648-658) that could not be unambiguously modelled in the maps, the final model encompasses the entire hTRPV4 sequence (residues 148-786). The non-protein density was obtained by simulating a 6Å map from the model of hTRPV4 using the molmap command integrated in the UCSF Chimera software and subtracting this theoretical apo-density form the experimental ligand-bound cryo-EM density [39].

### Docking

The GlideEM package (Schrodinger Maestro version 2019-4) was used for initial 4α-PDD docking calculations [48]. During preparation of the coordinates, residues not resolved in the final hTRPV4 model were generated by homology modelling using the xTRPV4 model as a template [25]. All GlideEM calculations were performed with default parameters and assuming an approximate EM map resolution of 4 Å using default values. The initial sampling phase generated a large number of candidate poses that were then filtered into the five top scoring poses for the refinement phase and manual inspection. These five top poses displayed a similar placement of the 4α-PDD phorbol group and differed only in the detailed position of the acyl chains pointing toward the membrane. Real-space refinement was then performed using the software Phenix [49] and utilizing the state-of-the-art OPLS3e/VSGB2.1 force field [50]. Following refinement, the top scoring pose was selected based on glide scores, visual inspection of the correlation with the non-protein electron density and manual inspection of the chemical interactions between ligand and the protein environment.

### Homology Modelling

A closed conformation tetrameric homology model of hTRPV4 was generated using the cryo-EM structure of xTRPV4 as a template (PDB entry 6BBJ [25]). Model building was performed with the MODELLER software [51] within Discovery Studio (BIOVIA; Dassault Systemes) using standard parameters and a high optimization level during sampling in the simulated annealing step. Resulting models were ranked according to their probability density function (PDF) energy, derived from spatial restraints when building the initial models. The model with the lowest PDF energy was used for further analysis.

### Whole-cell voltage clamp measurements on high five cells

In advance to transfection, high five-cells were transferred into culture medium without antibiotics and reduced proportion of FBS (1.5 %). 105 cells were plated onto glass cover slips coated with concanavalin-A (400 μg/ml) and laminin (4 μg/ml). Transfection required two separate mixtures, mixture A: 3.3 μL cellfectin (Invitrogen, 10362-100) and 42 μL medium (Express Five^®^ SFM + 90 mL 100x GlutaMax™, gibco, 10486-025), with an incubation time of 30 min; and mixture B: 1 μg plasmid (encoding either full-length and truncated human TRPV4 constructs with a C-terminal eGFP tag in a modified pXINSECT-DEST38 vector (Invitrogen), 2 μL Reagent plus (Invitrogen, 11514-015) and 42 μL medium (Express Five^®^ SFM + 90 mL 100x GlutaMax™) with an incubation time of 5 min. Mixture A and B were combined and, after an incubation of 15 – 30 min at rt, added to the cells. 4 h after transfection the old medium was discarded and the cells were incubated at 27 °C with medium containing 10 % FBS until they were ready for measurements (24-48h).

For whole cell voltage-clamp measurements high five cells transient expressing the human TRPV4 channel were plated onto glass cover slips previously coated with concanavalin-A (400 μg/ml) and laminin (4 μg/ml). The cells were kept at 27 °C. Electrophysiological recordings were done with the whole-cell voltage technique as described elsewhere [52]. The external bath contained Ringer’s solution: 150 mM NaCl, 4mM KCl, 2mM MgCl_2_, 2mM CaCl_2_, 10mM HEPES (pH 7.4 adjusted with NaOH). The (internal) pipette solution contained 120mM KF, 30mM KCl, 10mM K-EGTA, 1mM CaCl_2_, 10mM HEPES (pH 7.4 adjusted with KOH). Compounds were applied to the cells using the U-tube reversed flow technique [53].

The test compounds were freshly dissolved as a 10 mM stock solution in DMSO and diluted to the required concentrations in Ringer’s solution before an experiment. Currents were measured with the L/M-EPC 7 patch clamp amplifier (List, Darmstadt, Germany) and HEKA EPC 10 patch clamp amplifier (HEKA, Ludwigshafen, Germany). Current records were low-pass Bessel filtered at 1 kHz (EPC7) and 3 kHz (HEKA EPC 10) and digitized at 3 kHz sample rate (EPC7) and 10 kHz (HEKA EPC10).

### Two-electrode voltage clamp measurements on *Xenopus oocytes* expressing TRPV4 variants in oocytes

*In-vitro* transcription was performed with a linearized plasmid as DNA-template. Plasmid-DNA was cleaved with a suitable restriction enzyme, by following the instructor’s manual. The obtained linear DNA strand was then transcripted to RNA (*in-vitro* transcription-kit: mMESSAGE mMACHINE™ T7 Transcription Kit, ThermoFischer, Art.-No.: AM1344). The RNA concentration was measured and diluted to 200 ng/μL (nuclease free water).

*Xenopus laevis* oocytes (EcoCyte, GER) were incubated for 4 h at 19°C in Bath’s-solution (96 mM NaCl, 2 mM KCL, 1.8 mM CaCl_2_, 0.82 mM MgCl_2_, 50 μg/mL gentamycin, 1 μM ruthenium red; pH 7.6). The oocytes were transfected with ssRNA using RoboInject (multichannel systems, GER). For heterologous expression, 50 nL of the appropriate 200ng/μL RNA-solution were injected. After 48 h (full-length) and 72 h (truncated) hTRPV4 incubation at 19°C, the oocytes were measured.

Electrophysiological oocyte experiments were performed in the two-electrode voltage clamp mode. All oocytes were measured with the RoboCyte-Setup (1 & 2) of multichannel systems (GER). The bath solution contained 96 mM NaCl, 2 mM KCL, 0.3 mM CaCl_2_, 1 mM MgCl_2_, 5 mM HEPES and was adjusted to pH 7.6, whereas the pipette solution contained 1 M KCl and 1.5 M KAc with pH 7.2. All measurements were performed with corrected liquid junction potential at room temperature. In the measurements, cells were clamped to a resting potential of 60 mV and exposed to different concentrations of GSK1016790A and 4α-PDD.

### Characterization of TRPV4 mutants using calcium flux assays

Plasmids containing hTRPV4 (Gene ID: 59341) in a PiggyBac vector were obtained by custom synthesis from DNA Cloning Services (Hamburg). Point mutants were introduced at DNA Cloning Services (Hamburg) using site-directed mutagenesis. Chinese hamster ovary-K1 cells (CHO-K1) were maintained in DMEM/F12 supplemented with 10% FCS, 1.3% HEPES, 1% sodium pyruvate, 1% sodium bicarbonate, 1% P/S at 37°C and 5% CO_2_. Cells for functional characterization of agonist activity were created by transfection of CHO-K1 cells with TRPV4 and selection using G418 for at least one week. Wildtype TRPV4 cells were subcloned to yield a stable monoclonal cell via limited dilution [54]. All other mutants were used as stable clonal pool.

For calcium flux measurements, cells were seeded one day prior to the measurement in DMEM/F12 supplemented with 2% FCS, 1.3% HEPES, 1% sodium pyruvate, 1% sodium bicarbonate, 1% P/S at 37°C and 5% CO_2_ in 384 well plates (Greiner F-Bottom, μCLEAR, TC treated) at a concentration of 5000 cells per well. To measure intracellular calcium, cells were incubated with a Tyrode solution containing 1.2μM Fluo-8, 0.05% Pluronic acid, 42mM Probenicid and 166 μg/ml Brilliant Black one hour before the measurement at 33°C and 5% CO_2_. Before each measurement, agonists were dissolved and diluted in DMSO to yield half log concentration curves. Calcium flux measurements were performed on a 384-well FLIPR Tetra (Molecular Devices) with fluorescence excitation at 480nm and fluorescence detection at 520nm. After pre-integration for 5s (1 frame per second), agonists were added as a 4x concentrated solution in Tyrode containing 0.01% BSA and 166μg/ml Brilliant Black. Kinetics were recorded for 190s. During each experiment, each agonist concentration was measured as a quadruplicate. The calcium signal was extracted after 190s, the data normalized to the response to the specific TRPV4 agonist GSK1016790A at 25μM and dose response curves were fitted using in-house software. Each experimental condition yielding a calcium flux response was repeated at least 3 times to obtain an average EC_50_ and standard deviation. Non-responding mutant variants were repeated twice. EC_50_ was defined as >25 μM, if no fit was possible.

## ADDITIONAL INFORMATION

### Data availability

The cryo-EM map and atomic model were deposited in the Electron Microscopy Data Bank and the Protein Data Bank (PDB) under the accession codes EMD-11690 and PDB 7AA5, respectively.

### Author Contributions

M.B, A.B., U.E., H.S., M.H. and S.J.H. conceived the project. M.B. and A.K.C.U., with guidance from N.B. and V.P., cloned and established biochemical conditions for protein preparation. M.B and R.A. performed cryo-EM grid freezing, data collection and data processing M.C. assisted with cryo-EM grid freezing and supported EM data collection. M.B. and D.B. built the hTRPV4 atomic model. D.B. built the 4α-PDD binding model. A.B. and U. E-K. performed and analyzed electrophysiological characterization measurements. D.G. performed and analyzed calcium flux mutant characterization measurements. U.E. built the closed conformation homology model and performed detailed structural comparisons. M.H and S.J.H. coordinated the project. S.J.H. wrote the manuscript with input from all authors.

## ACKNOWLEDGMENTS

The expert technical assistance of Sabine Daemmig and Laura Bohmann is gratefully acknowledged. We also thank Nicolas Werbeck, Cora Sholten and Anke Mueller-Fahrnow for helpful discussions and support. This work was in part supported by the Swiss National Science Foundation, grant NCCR TransCure.

---

**Extended Data Table 1:**
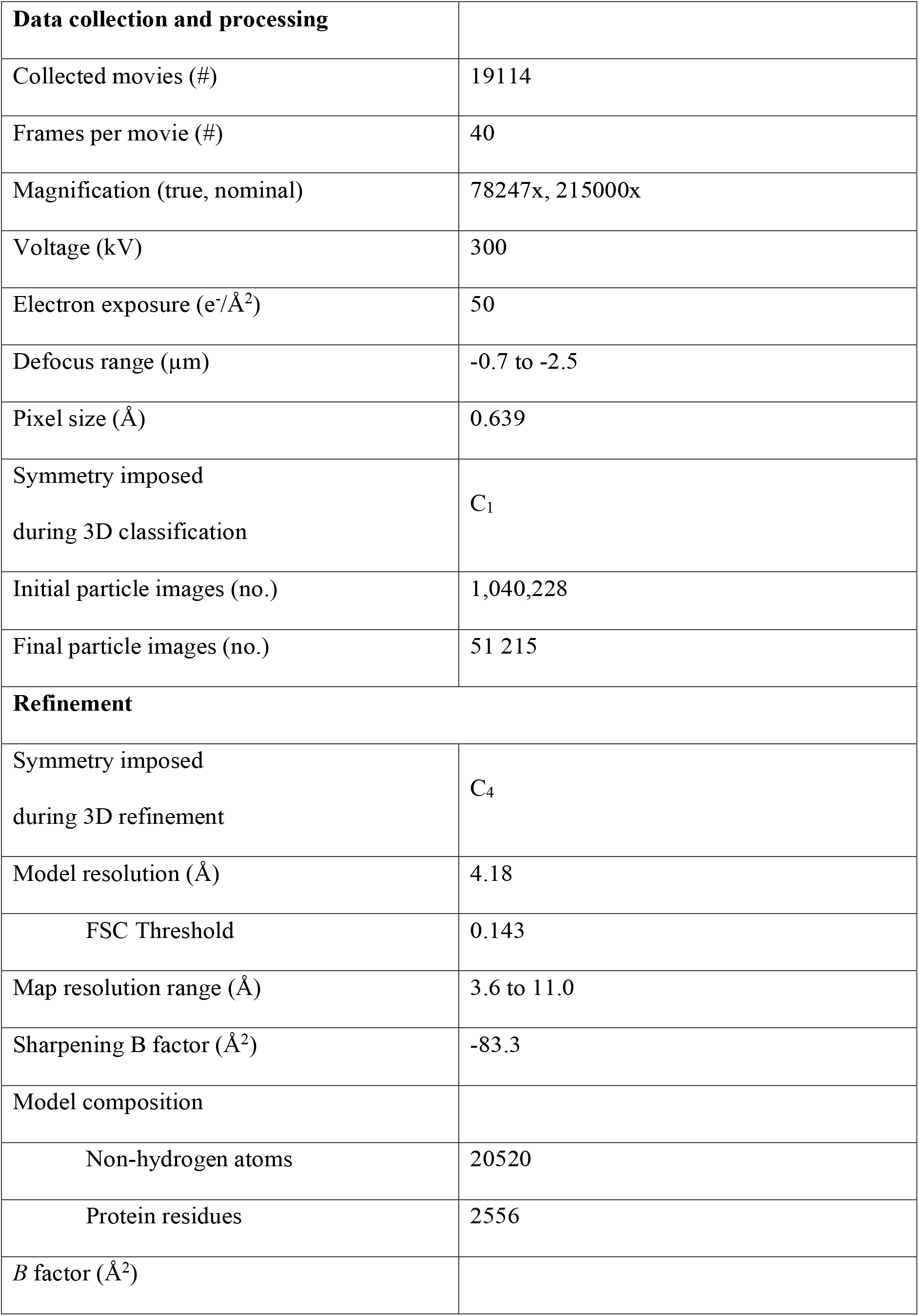
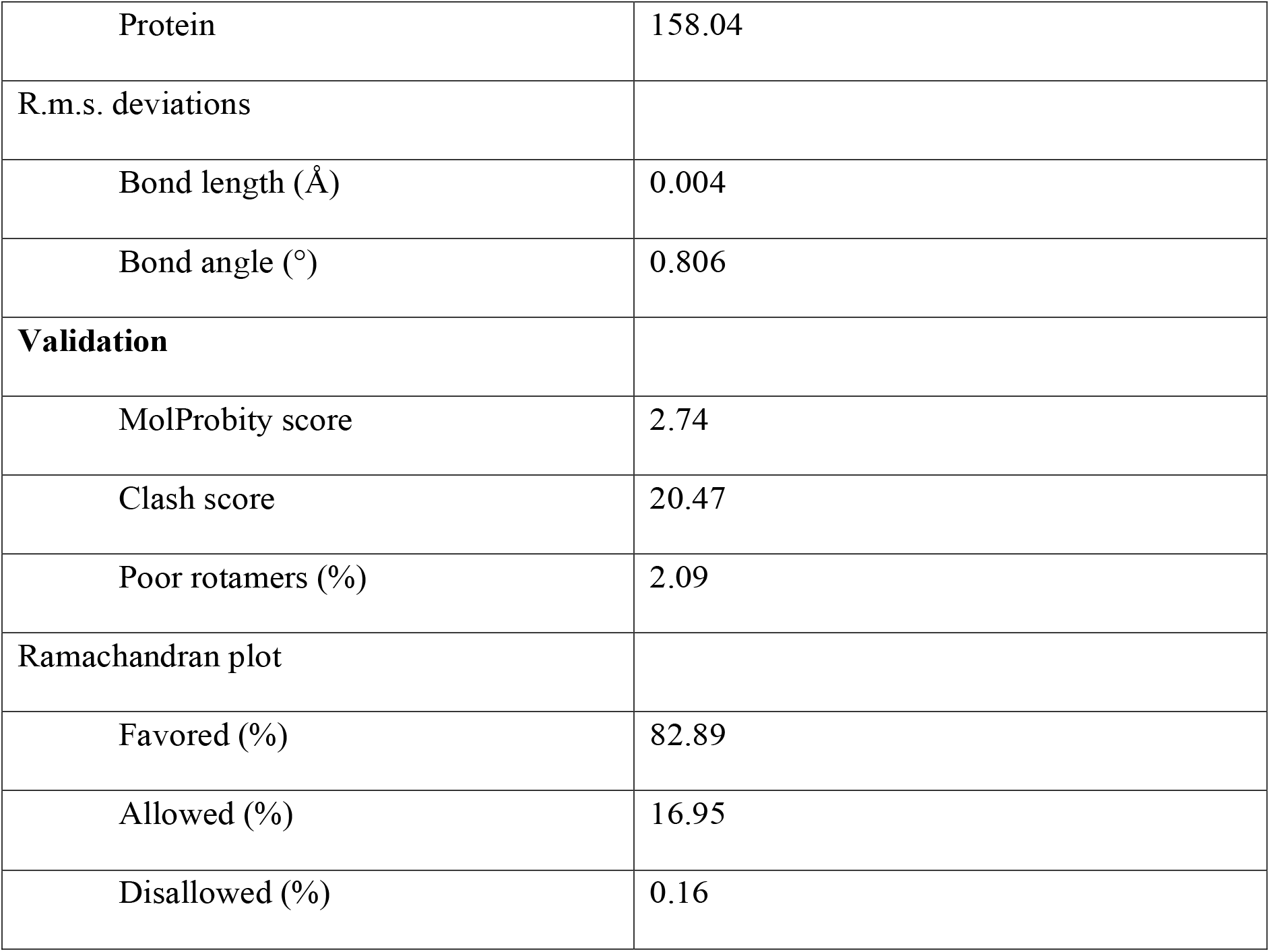
Cryo-EM data collection, refinement and validation

